# Expansion microscopy allows quantitative characterisation of structural organisation of platelet aggregates

**DOI:** 10.1101/2024.09.16.613233

**Authors:** Emma L. Faulkner, Jeremy A. Pike, Evelyn Garlick, Robert K. Neely, Iain B. Styles, Stephen P. Watson, Natalie S. Poulter, Steven G. Thomas

## Abstract

Current microscopy approaches applied to platelet aggregates in both haemostatic and thrombotic settings indicate their structure has important implications in efficient haemostasis and in clinical treatment of thrombosis. However, current fluorescence microscopy approaches are not amenable to volumetric imaging of platelet aggregate structures. This is largely due to the small size of individual platelets and the tight packing of platelets within aggregates, resulting in optical opacity.

Here we demonstrate that expansion microscopy, applied to platelet aggregates, can reveal multi-scale information about the structure of platelet aggregates. We produced volumetric images at nanoscale resolution of >700 platelet aggregates under normal and perturbed conditions, stained for cytoskeletal and membrane components. We demonstrate our custom analysis workflow provides quantitative description of platelet numbers, volumes and morphology within entire platelet aggregates. Additionally, we quantitatively describe subcellular organisation of F-actin. By comparing these measurements following treatment with the actin inhibitors, cytochalasin D and latrunculin A, we can robustly detect structural disruptions in platelet aggregates. Together these data provide a workflow to qualitatively and quantitatively describe the architecture of platelet aggregates at a range of scales (whole aggregates down to sub-cellular features within individual platelets).

## 2 Introduction

The function of platelets in haemostasis and thrombosis is principally reliant on their ability to rapidly activate and adhere at sites where blood vessels are perturbed due to vascular injury, damage or inflammation. The cytoskeleton plays a critical role in platelet function, maintaining the discoid shape of circulating resting platelets and driving dynamic rearrangements in response to activation [1]. The actin cytoskeleton is particularly important for platelet shape change and has been shown, *in vitro*, to form specific F-actin structures [2]. Actin nodules form in the early stages of platelet adhesion and spreading on different ligands, with F-actin stress fibres formed in the later stages [3, 4]. Whether these structures form in platelet aggregates or thrombi is not yet known. Actin and actin binding proteins also play roles in regulating platelet receptors, for example integrins [5] and (hem)ITAM receptors [6, 7]. Actin-myosin is also important for thrombus contraction and stability [8–10]. Thrombus formation is a multistep process, initiated by multiple factors including interaction of the platelet GPIb-V-IX complex with immobile von Willebrand factor and binding of GPVI to collagen [11]. This drives a shift of platelet integrins to their higher affinity state, supporting strong adhesion and platelet aggregation [12]. Production of thrombin and other soluble mediators sustain the platelet aggregate and drive deposition of fibrin to stabilise the thrombus. Due to the complexity of interactions required for thrombus formation, the resulting thrombi are heterogeneous structures comprised of multiple components (e.g. fibrin, erythrocytes, platelets, leukocytes and neutrophil extracellular traps (NETs)) [13–15].

The process of platelet aggregation has been studied extensively, revealing that thrombus structure is critical to correct haemostatic function. Intravital and electron microscopy (EM) approaches have revealed *in vivo* haemostatic thrombi adopt a hierarchical structure, with a core of highly activated, tightly packed platelets overlaid by a shell of less activated, loosely packed platelets [16, 17]. This architecture is critical to stabilizing and limiting overall thrombus growth.

The conditions that result in arterial thrombosis, where platelet-rich thrombi form in response to atherosclerotic plaque rupture/erosion, and immunothrombosis, where thrombi are initiated by an immune response [18] are different to their haemostatic counterparts and results in different thrombus architecture. It has also been shown that thrombus composition differs depending on factors such as vascular bed, thrombotic trigger and blood flow [13, 19]. Understanding the composition and architecture of thrombi has implications in characterising disease progression and in improving treatments and clinical outcomes [13, 20, 21]. With these observations, facilitated by advancing microscopy techniques, there has been a shift from considering platelet aggregation from an agonist centric point of view to a more holistic overview of platelet aggregate architecture.

Imaging platelet aggregates is complicated by (i) the small size of platelets (approx. 2 μm diameter), (ii) the tight packing of platelets, other cell types and fibrous components in the thrombi and, (iii) the relative optical opacity of platelet aggregates. As a result, resolving sub-cellular features within individual platelets at nanoscale resolution is challenging, and three-dimensional (3D) volumetric imaging of whole thrombi is hampered by optical inaccessibility. Here we demonstrate that expansion microscopy (ExM) can overcome these challenges to facilitate qualitative and quantitative imaging of the structural organisation of platelet-rich thrombi. This is achieved by reducing the spatial density of labelled features resulting in increased resolution, and by optically clearing samples, thus facilitating volumetric imaging with relative ease [22–24].

## 3 Method and Materials

### 3.1 Antibodies and Reagents

All chemicals, unless otherwise stated, are from Sigma or ThermoFisher. Horm collagen from Nycomed. Primary Antibodies: anti-human CD41 antibody (Agilent, F708801-2) and alpha tubulin antibody (Sigma Aldrich, Clone DM1A, sc-32293). Secondary antibodies: goat anti-mouse IgG conjugated to CF633 dye (Biotium, 20120). Actin ExM probe conjugated with 561 dye was purchased from Chrometra.

### 3.2 Blood collection

Blood collection from healthy human volunteers was via venepuncture into 0.1 (v/v) volume of sterile 129 mM trisodium citrate under ethics granted by University of Birmingham internal ethical review (ERN-11-0175). Citrated blood samples were recalcified with 3.75 mM MgCl_2_ and 7.5 mM CaCl_2_ in the presence of PPACK (40 μM) prior to the flow experiment, as described in Jooss *et al* [25].

### 3.3 Platelet preparation and spreading

Washed platelets were prepared as previously described [26] from acid-citrate-dextrose-anticoagulated blood from healthy volunteers and resuspended at 2 x 10^7^ platelets mL^−1^ in HEPES-Tyrodes buffer (134 mM NaCl, 0.34 mM Na_2_HPO_4_, 2.9 mM KCl, 12 mM NaHCO_3_, 20 mM HEPES, 5 mM glucose, pH 7.3). Round coverslips (13 mm, #1.5) were incubated with fibrinogen (100 µg mL^−1^) overnight at 4°C. Coverslips were blocked in 5 mg mL^−1^ bovine serum albumin (BSA) for 1 hour. Resuspended platelets were incubated with the coverslips for 45 minutes at 37°C. Spread platelets were fixed in 10% (v/v) neutral buffered formalin and washed three times with PBS.

### 3.4 Coating coverslips

Glass coverslips (22×50 mm #1.5) were washed in ethanol followed by saline and water then allowed to dry overnight. Collagen was diluted in the supplier provided diluent. Collagen coating (1-2 µL) was applied to each coverslip at a concentration of 100 µg mL^−1^. Collagen was thoroughly vortexed prior to dilution and coating. Coverslips were incubated in a humidified chamber at 4°C overnight then washed twice with PBS before being blocked with HEPES buffer pH 7.45 (136 mM NaCl, 10 mM HEPES, 2.7 mM KCl, 2 mM MgCl_2_) supplemented with 0.1 % (w/v) glucose and 1 % (w/v) bovine serum albumin (BSA) for 45 minutes at room temperature. If stored, coverslips were immersed in HEPES buffer pH 7.45 (136 mM NaCl, 10 mM HEPES, 2.7 mM KCl, 2 mM MgCl_2_) and stored at room temperature until required.

### 3.5 Flow chamber device and perfusion protocol

Collagen-coated coverslips were mounted onto the parallel plate flow chamber [27] and pre-rinsed with HEPES buffer pH 7.45 to remove any air bubbles. Anti-coagulated whole-blood samples (500 μl) were perfused through the flow chamber for 3 minutes at 1,000 s^−1^ to allow platelet aggregate formation on collagen. Flow was monitored using an EVOS microscope to confirm laminar flow and absence of air bubbles. The flow chamber was then perfused with HEPES buffer pH 7.45 for 2 minutes. Following 2 minutes of stasis the platelet aggregates were then washed with HEPES buffer at 1000 s^−1^. The flow chamber was then deconstructed and platelet aggregates were fixed in 10 % (v/v) neutral buffered formalin (CellPath, BAF-6000-08A) for 10 minutes at room temperature. Coverslips were then washed with PBS and stored in PBS prior to immunofluorescent staining.

### 3.6 Treatment of pre-formed platelet aggregates with Cytochalasin D (CytD) and Latruncullin A (LatA)

Following perfusion of anticoagulated blood through the flow chamber for 3 minutes at 1000 s^−1^, actin inhibitors, diluted to the stated concentrations in HEPES buffer pH 7.45, were perfused through the chamber at 1000 s^−1^ for 10 minutes. Following 2 minutes of stasis, platelet aggregates were washed with HEPES buffer and fixed, as stated above.

### 3.7 Immunofluorescence staining

Spread platelets and platelet aggregates were subject to immunofluorescent staining prior to standard microscopy using a widefield microscope or were prepared using ExM methodology. A well was drawn around platelet aggregates with a hydrophobic pen on each coverslip to minimise reagent use throughout staining and to simplify the gelation and digestion steps required for the ExM preparation. Platelets were permeabilised with 0.1 % (v/v) Triton X-100 in PBS for 5 minutes at room temperature. After blocking with 1 % (w/v) bovine serum albumin (BSA) and 2 % (v/v) goat serum in PBS for 30 min, platelets were incubated with primary antibodies diluted in blocking buffer at the following dilutions: anti-human CD41 monoclonal antibody (1:250) and alpha tubulin antibody (1:500). Platelets were washed three times with PBS and incubated with goat anti-mouse IgG-CF633 (1:1000). Platelets were then washed three times with PBS. For spread platelets all antibody incubations were performed for 1 hour at room temperature. For platelet aggregates all antibody incubations were performed overnight at room temperature.

### 3.8 Sample Expansion

Acryloyl-X (6-((acryloyl)amino)hexanoic acid, succinimidyl ester) (AcX) (A20770, ThermoFisher Scientific) was re-suspended in anhydrous DMSO with a final concentration of 10 mg mL^−1^. This was then aliquoted and stored in a frozen desiccated environment for up to 2 months. Platelet aggregates were incubated with the AcX at a final concentration of 0.1 mg mL^−1^ in PBS for >6 hours at room temperature. Specimens were washed with PBS and then labelled with Actin ExM (Chrometra) at 1 unit/coverslip for 1 hour. Anchoring with AcX prior to Actin ExM labelling was essential to ensure undistorted retention of the Actin ExM probe post-ExM. Gelation was performed immediately after 3 washes with PBS. Monomer solution (1xPBS, 2 M NaCl, 8.625 % (w/w) Sodium acrylate (97 %, 744-81-3, Sigma Aldrich), 2.5 % (w/w) acrylamide (79-06-1 Sigma Aldrich), 0.15 % (w/w) N,N’-methylenebisacrylamide (110-26-9, Sigma Aldrich)) was mixed, frozen in aliquots and thawed prior to use. Concentrated stocks of ammonium persulfate (APS) (7727-54-0, Sigma Aldrich) and tetramethylethylenediamine (TEMED) (110-18-9, Sigma Aldrich) at 10 % (w/w) in water were diluted into the monomer solution to final concentrations of 0.2 % (w/w) on ice prior to gelation, with the initiator (APS) added last. The gelation solution (120 µL) was added to the well formed by a hydrophobic pen on each coverslip. For spread platelets, round coverslips were inverted onto a droplet (90 µl) of gelation solution on a parafilm-coated slide. Gelation was allowed to proceed at 37°C for 2 hours in a humidified chamber. Gels were removed from the coverslips and immersed in digestion buffer (1xTAE, 0.5 % (v/v) Triton X-100, 0.8 M guanidine HCl) containing 8 units mL^−1^ of Proteinase K (P8107S, New England Biolabs Inc.). Gels were digested at room temperature overnight. The gels were removed from the digestion buffer and placed in 50 mL of water to expand. Water was exchanged every 30 minutes until expansion was complete (typically 3-4 exchanges).

### 3.9 Expanded specimen handling for imaging

For widefield imaging, expanded gels were cut to fit in imaging dishes with a glass coverslips of 35 mm diameter (81218-200, Ibidi). Excess water was removed and gels were embedded in 2 % (w/v) agarose to limit gel movement during image acquisition. For Selective Plane Illumination Microscope (SPIM) imaging, gels were cut to fit the SPIM holder and placed cell-side up in it. 2 % (w/v) agarose was pipetted into the holder until the bottom of the holder was covered, taking care not to get agarose in the interface between the top of the ExM gel and the objective lens. Deionised water was then added to the SPIM holder containing the gel to fully immerse the gel for imaging.

### 3.10 Image acquisition

Samples were imaged using a Nikon N-SIM-S system (Ti-2 stand, Cairn TwinCam with 2 × Hamamatsu Flash 4 sCMOS cameras, Perfect Focus, Nikon laser bed 488, 561 and 647 nm excitation lasers, Nikon 100 × 1.49 NA TIRF Apo oil objective or Nikon 100 × 1.35 NA Silicone oil objective. NIS-elements software was used to control the system). General imaging and screening of thrombi was performed using a Zeiss Axio Observer 7 inverted epifluorescence microscope (Definite Focus 2, Colibri 7 LED illumination source, Hamamatsu Flash 4 V2 sCMOS camera, Filter sets 38, 45 and 50 for Alexa488, Alexa561 and Alexa647, respectively. Acquisition was performed using Zen 2.3 Pro software). To generate 3D volumes for quantitation of platelets within thrombi a SPIM microscope was used (Marianas diSPIM system with ASI RAMM microscope frame with twin Hamamatsu ORCA-Flash4.0 v3 sCMOS cameras, 488, 561, 640 laser lines via Shemrock 440/521/607/700 Brightline quad bandpass filter and twin Nikon CFI NIR Apo 40x 0.8 NA water dipping objectives. Slidebook 6 software was used to control the system. All images were acquired using stage scanning).

### 3.11 Image processing

Where stated, post-ExM image data was deconvolved using Huygens professional version 19.04 (Scientific Volume imaging, the Netherlands, http://svi.nl). A theoretical point spread function (PSF) was generated based on the microscope parameters and images were deconvolved using a classical maximum likelihood estimation (CMLE), a non-linear iterative restoration method which optimises the likelihood the objects in the estimated image are correctly localised based on the image and the PSF. Parameters for deconvolution were tested on example data sets for each experiment to determine optimal values, and these deconvolution templates were used for subsequent image processing and experimental repeats. Images acquired on the Marianas diSPIM were deskewed and interpolated prior to image reconstruction and analysis. Images were stored as Slidebook file format (*.sld)

### 3.12 Segmentation of individual platelets in platelet aggregates using CellPose segmentation algorithm

For segmentation of platelets within platelet aggregates, the deskewed diSPIM data were sorted into directories according to imaging conditions. From each of the acquired .sld documents, each series was exported as a 16 bit TIFF file using a built-in FIJI macro (batch_convert.ijm) [28]. A Python script (predict_regions.py) was applied which allowed identification of individual aggregates as regions of interest (ROIs) using an Otsu threshold and connected component analysis. The identified aggregates were cropped by a bounding box determined from the connected component analysis. Cell segmentation was performed on the cropped regions using the pretrained “cyto” Cellpose model with subsequent stitching of 2D results to produce 3D segmentations [19]. A second Python script (post_process.py) opened each of the images produced by predict_regions.py and used the cellpose segmentations and the Otsu masks to compute a range of measurements for each identified cell in the aggregate using built in algorithms from the scikit-image Python package [29]. Measurements, which included volume, boundary pixels, intensity, and calculations of the membrane and cytoplasm volumes, were saved as a .csv file from which data was be subsequently analysed using Microsoft Excel 365 and Graphpad Prism 10 as detailed in figure legends. Visualisation and inspection of analysis results for a specific aggregate using Napari was performed with the expansion_visualisation.ipynb Jupyter notebook [30]. All Python scripts and the notebook can be found on the GitLab repository at https://gitlab.bham.ac.uk/thomass-expansion-microscopy/cellpose-expansion-analysis. The full data sets generated from the 5 different healthy volunteer donors and those following treatment with actin inhibitors are available as a data supplements and include the ability to filter data by donor, label, etc. (Data Supplement 1 & 2). Identified platelet aggregates and individual platelets for each experimental condition are summarised in Supplementary Table 1.

## 4 Results

### 4.1 Validation of ExM for platelet aggregates

We hypothesised ExM preparation would circumvent existing limitations in obtaining 3D images of platelet aggregates with nanoscale resolution of subcellular components. To investigate this, we applied a four-fold ExM preparation [31] to individual spread platelets and platelet aggregates (illustrated in Figure 1A). Spread platelets were stained for α-tubulin and imaged pre-ExM and post-ExM. Post-ExM images were also deconvolved to increase contrast and resolution (Figure 1B). Following ExM preparation (Post-ExM), imaged platelets occupied an increased surface area and an appreciable increase in resolution as compared to related samples pre-ExM. A lateral effective resolution of 75 nm was determined by measuring full width half maximum (FWHM) of line profiles of microtubules in deconvolved post-ExM samples (Figure 1C), a resolution improvement consistent with the fourfold macroscale expansion of the samples. Images were acquired pre-ExM (Figure 1D) and post-ExM (Figure 1E) of platelet aggregates stained for the platelet integrin component CD41 and F-actin. We observed an increase in aggregate size and in lateral and axial resolution which was further enhanced by the optical clearing properties of the ExM preparation.

**Figure 1.**
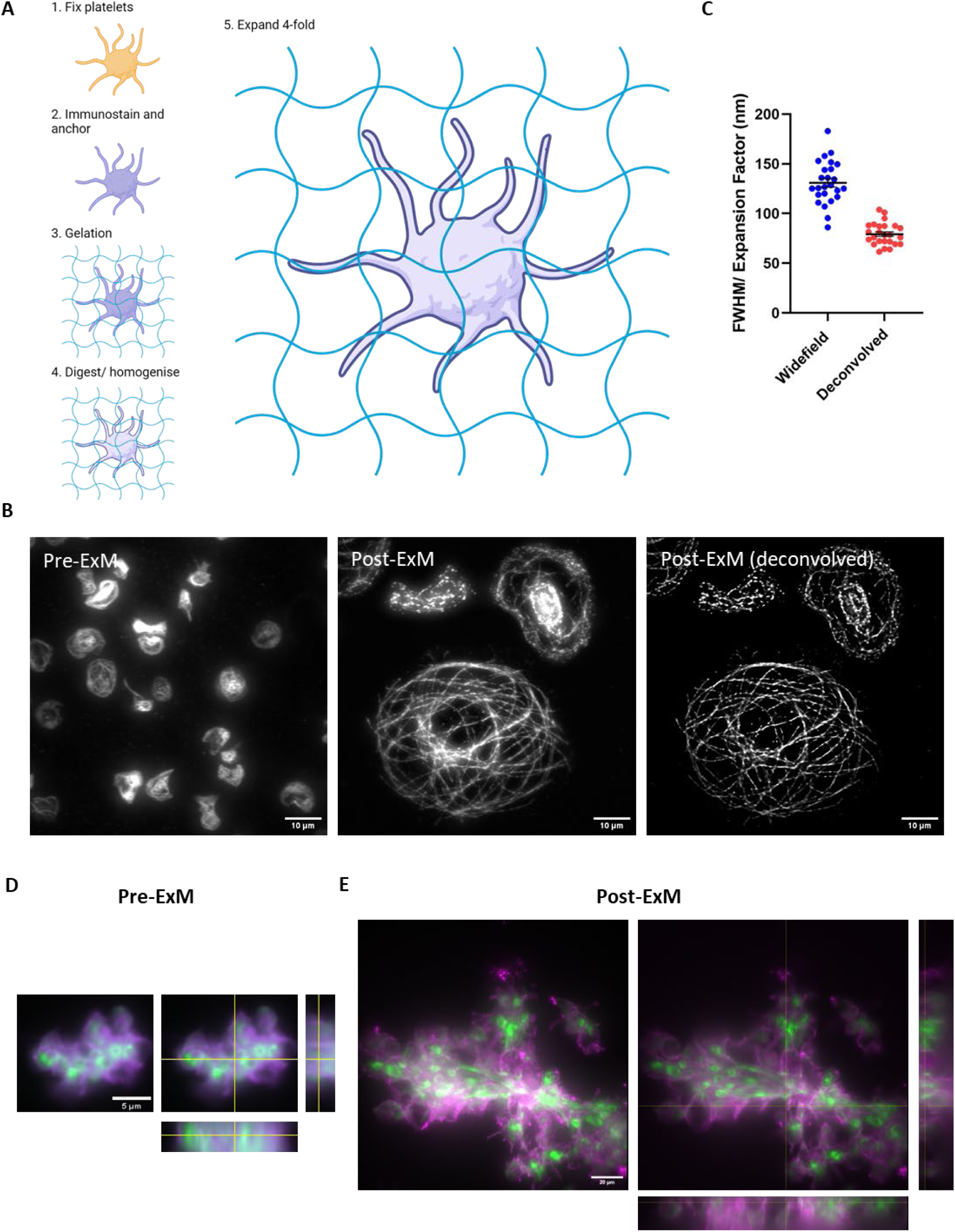
Expansion microscopy preparation enables nanoscale resolution of subcellular platelet features and increases imaging depth and volume. A) Schematic of ExM preparation of platelets. Platelets are fixed, immunostained and then anchored with acryloyl-x. Platelets are embedded in a polyelectrolyte gel in which labels are bound to the gel via acryloyl-x. Samples are then homogenised using proteinase K and finally expanded by addition of water, resulting in a four-fold isotropically expanded sample. B) Washed platelets were spread onto fibrinogen coated coverslips and immunostained for tubulin. Images were either acquired pre- or post-ExM on an epifluorescence microscope. Post-ExM images were deconvolved using Huygens software. Scale bars 10 µm. C) Effective resolution of expanded samples were determined by measuring the full width half maximum (FWHM) of microtubules post-ExM before and after deconvolution. Mean ± SEM. Measured values were corrected for the expansion factor determined by measuring the gel diameter. D) Platelet aggregates were fixed and stained for CD41 (magenta) and phalloidin (green). Pre-ExM and post-ExM images were acquired on an epifluorescence microscope. Z stacks are represented with orthogonal projections for each dimension. Scale bars - pre-ExM 5 µm and post-ExM 20 µm. Images in D representative of five repeats.

### 4.2 Investigating 3D organisation of membrane and cytoskeletal components in platelet aggregates

We characterised the structural organisation of platelet aggregates formed on collagen from multiple healthy donors. For each healthy donor, platelet aggregates were stained for CD41, and the cytoskeletal components tubulin and F-actin. Stained platelet aggregates were prepared for ExM and post-ExM images were acquired either on an epifluorescence widefield microscope (Supplementary Figure 1) or a selective plane illumination microscope (SPIM) (Figure 2). Examples of 3D image acquisitions are provided (Supplementary Videos 1-4).

**Figure 2.**
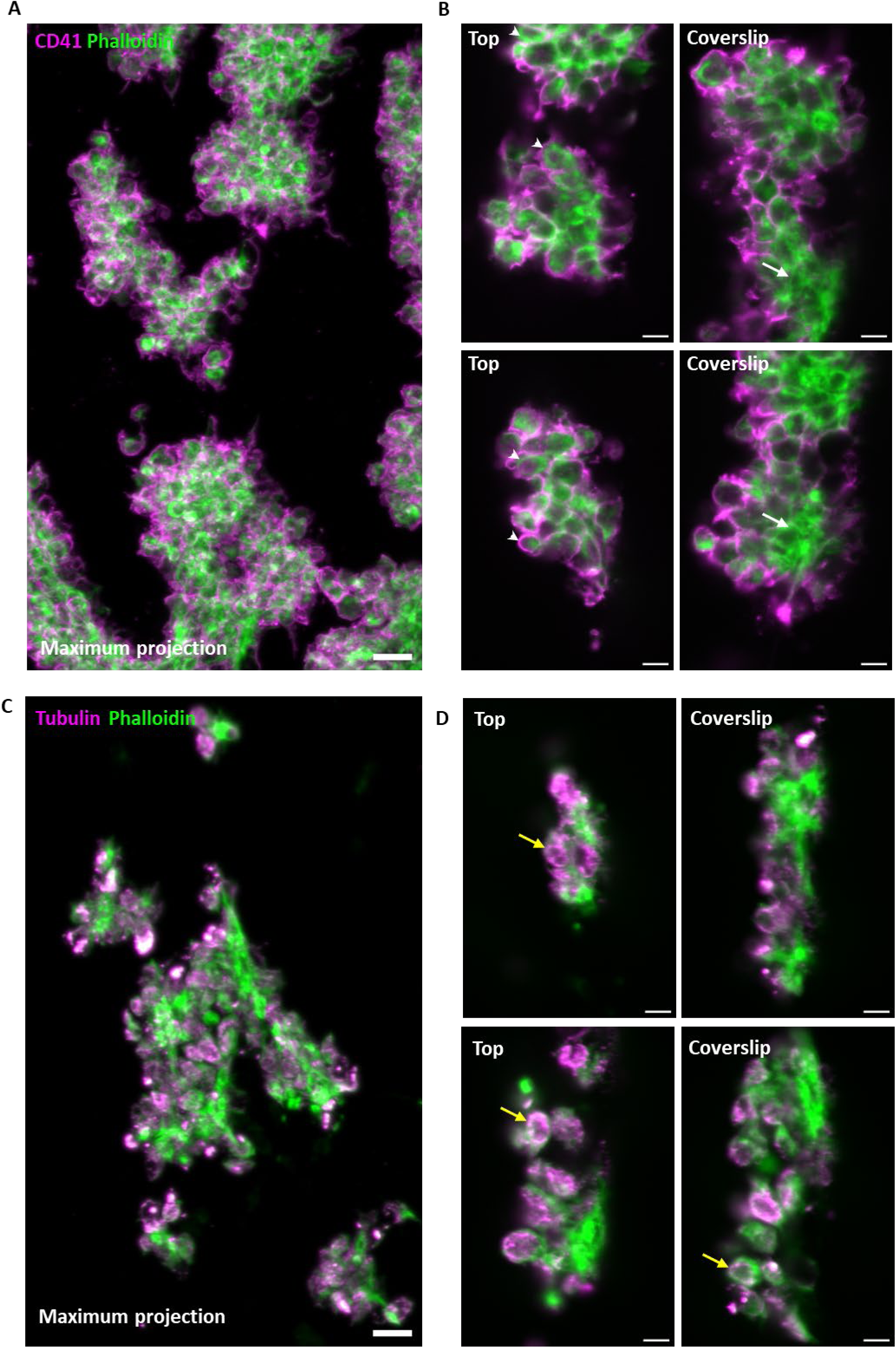
Expansion microscopy reveals heterogeneous distribution of membrane and cytoskeletal components in platelet aggregates. Platelet aggregates were prepared in parallel plate flow chambers under arterial shear (1000 s^−1^) on collagen-coated coverslips. Aggregates were fixed and immunostained for A & B) CD41 (magenta) and F-actin (green) or C & D) α-tubulin (magenta) and F-actin (green). Images of the aggregates were acquired on a SPIM. A & C) maximum projections of aggregates. B & D) Example focal planes taken from different aggregate at regions near the coverslip or at the top of the aggregate. White arrows – actin networks, white arrowheads – cortical actin, yellow arrows – tubulin rings. Scale bars. A & C – 20 µm, B & D – 10 µm. n=5.

We observed structural heterogeneity in platelet aggregates labelled for CD41 and F-actin (Supplementary Figure 1A, Figure 2A-B). Platelets at the adhesion surface formed filopodia- and lamellipodia-like protrusions and possessed an actin meshwork (Figure 2B, white arrows) which appeared to cross through individual platelets. Platelets at the top of the aggregates adopted a rounder morphology with actin organised cortically at membrane proximal regions (Figure 2B, white arrowheads). In aggregates labelled for tubulin (Supplementary Figure 1B, Figure 2C-D), we observed tubulin ring-like structures in individual platelets (Figure 2D, yellow arrows) together with diffuse accumulations dispersed throughout the aggregate volume. Together, these images demonstrate that platelet aggregates can be expanded, and imaging allows details of the platelets and their cytoskeleton to be visualised.

### 4.3 Developing a workflow to quantitatively describe structure of platelet aggregates

To further interrogate aggregate images, we developed an analysis workflow to allow quantitative descriptions of platelet aggregates and the platelets within them (Figure 3A). This resulted in a data set of 11,471 platelets from 421 aggregates generated from 5 different donors. Briefly, an Otsu threshold was applied to each image series to identify individual aggregates which were then automatically cropped from the image. Individual platelets within the aggregate were segmented in 3D using the deep learning-based segmentation method, Cellpose [32] on the CD41 channel. Example outputs from this workflow are shown in Figure 3B. From the masks generated by Cellpose, we calculated features including overall aggregate volume, number of platelets within each aggregate, the volume of individual segmented platelets and morphology parameters.

**Figure 3.**
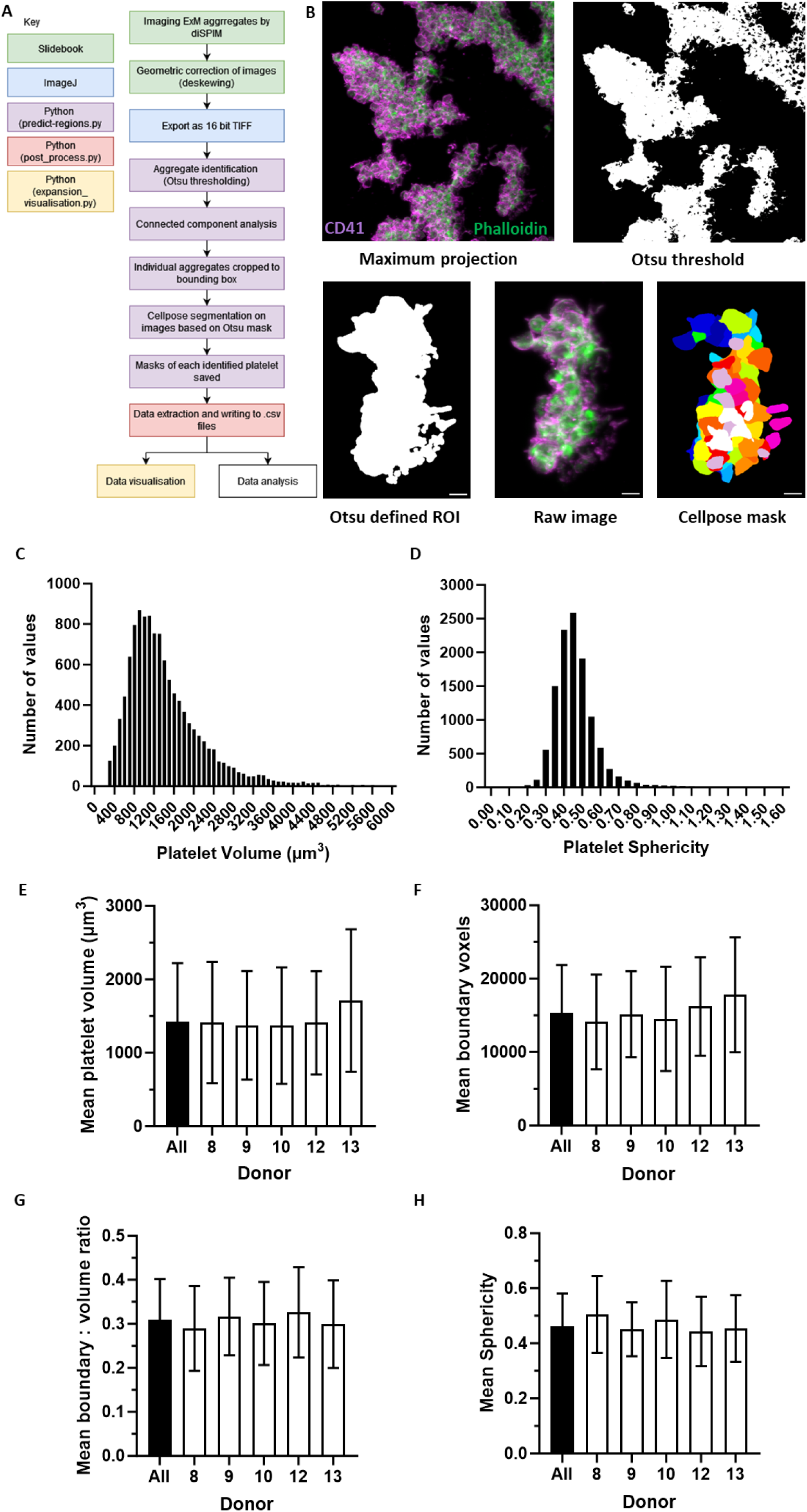
Analysis of platelet aggregates using light-sheet microscopy and custom analysis workflows. A) Flow chart of the image processing steps applied to expanded platelet aggregates images acquired on a selective plane illumination microscope (SPIM) with B) examples of deskewed maximum projections, application of the Otsu threshold, individual aggregate cropped to the boundary box, the raw data for comparison, and the identification of individual platelets within the aggregate based on Cellpose segmentation of the CD41 signal. C-H) Plots demonstrating donor to donor variation in a range of platelet parameters extracted from the custom image analysis workflows. E) platelet volume, F) Number of boundary voxels, G) boundary to volume ratio and H) platelet sphericity. Bars represent mean values ± SD. All donors, n = 11471 platelets; Donor 8, n = 1170 platelets; Donor 9, n = 6165 platelets; Donor 10, n = 2208 platelets; Donor 12, n = 453 platelets; Donor 13, n = 1475 platelets.

The average volume and sphericity of an expanded platelet within platelet aggregates was 1423 µm^3^ ± 797 and 0.46 ± 0.12 (mean ± SD) calculated from all platelets from 421 aggregates imaged (Figure 3C & D). The full data set of identified platelets is provided in Data Supplement 1. We compared platelet volumes, number of boundary voxels (a measure of the surface area of segmented platelets), the boundary to volume ratio, and platelet sphericity between the 5 blood donors used in our experiments. We observed some inter-donor variability but that all sit within the same overall range (Figure 3E-H). It is noticeable that donor number 13 has a larger mean platelet volume (1713 µm^3^ ± 970) than the other donors (mean = 1380 µm^3^ ± 759), and this is mirrored by a slightly larger mean number of boundary voxels for this donor (17799 ± 7833) compared to the other donors (mean = 14941 ± 6272). However, both the sphericity and the surface to volume ratio of the platelets from donor 13 were similar to the other donors. This demonstrates that this donor has slightly larger platelets than the other donors and was detected using this approach.

Additionally, the relationship between different parameters was assessed. There was a strong positive correlation between the number of boundary voxels and the platelet volume calculated on an individual platelet basis, and on an aggregate basis (individual platelets - R^2^ = 0.695, Supplementary Figure 2A; per aggregate – R^2^ = 0.5692, Supplementary Figure 2D). This demonstrates that the surface area of a platelet increases with volume and gives confidence that the segmentation of individual platelets is behaving as expected. Likewise, and as predicted, there is a strong positive correlation between the total aggregate volume and the number of platelets it contains (R^2^ = 0.9786, Supplementary Figure 2C). Conversely, there is no correlation between platelet sphericity and platelet volume (R^2^ = 0.0003, Supplementary Figure 2B) as the roundness of a platelet is not dependent on its volume. Together, this data highlights the power of our approach in allowing comprehensive qualitative and quantitative description of individual platelets from many aggregates at a level not previously achievable.

Given the observed heterogeneous organisation of F-actin in platelet aggregates across all donors, we sought to characterise the relative spatial distribution of actin to the platelet membrane using CD41 as a marker. Taking the Cellpose-determined boundary of each segmented platelet (Figure 4A), a membrane proximal region was defined by expanding the boundary by 3 voxels inwards (Figure 4B) with the remainder of the segmented volume defined as cytosolic (Figure 4C). The masks generated from these regions were used to characterise the actin organisation as being proximal to the membrane or dispersed through the cytosolic region. Data from the 5 healthy donors, indicated that on average there was a ∼1:1 ratio of membrane proximal to cytoplasmic actin distribution (Figure 4D & E).

**Figure 4.**
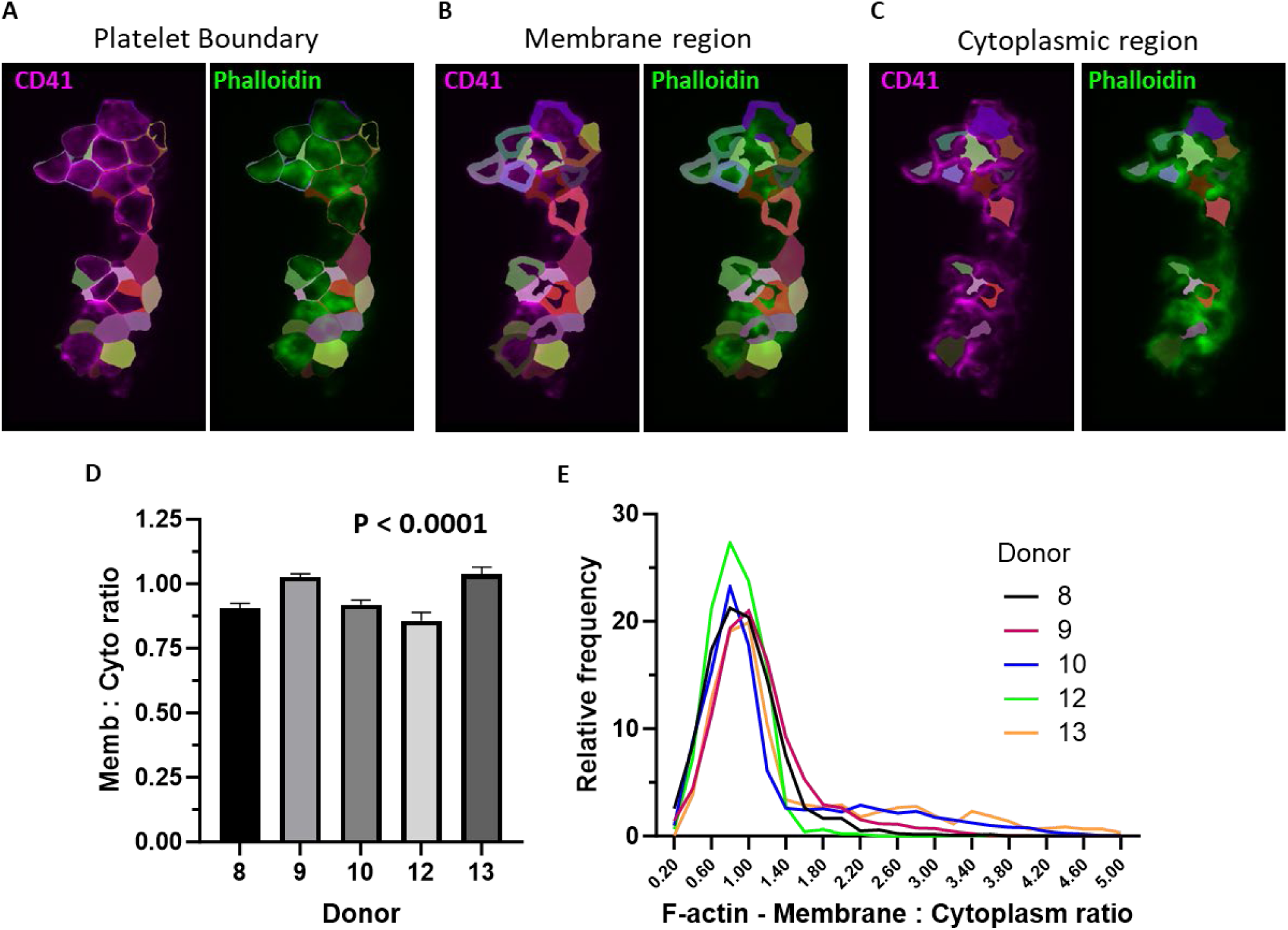
Classification of actin localisation in segmented platelets. A) Using the Cellpose mask, individual platelet boundaries within an aggregate were marked. B) The membrane region of the platelet was defined by expanding the platelet boundary inwards by 3 voxels and C) the remainder defined as the cytoplasmic region. Average voxel intensity in each region for the actin channel was then measured to describe the distribution of F-actin as either membrane or cytoplasmic. D) & E) Plots showing the median membrane : cytoplasmic ratio of F-actin from identified platelets from the 5 different donors. D) Median ± 95 % CI and E) frequency distribution. n = 5; 11,471 platelets.

### 4.4 Characterising structural impact of actin inhibitors on pre-formed platelet aggregates

It has been previously demonstrated that aggregate stabilisation is a dynamic process involving F-actin polymerisation; where F-actin dynamics are disrupted, aggregates become destabilised [33]. We hypothesised that by treating platelet aggregates with actin inhibitors we could induce structural disruptions which may be qualitatively and quantitatively assessed using the ExM approach. Here we applied cytochalasin D (CytD) or latrunculin A (LatA) to pre-formed aggregates to establish if this was the case.

We aimed to induce structural disruption in aggregates without inducing disaggregation, therefore pre-formed platelet aggregates were treated for 10 minutes with CytD and LatA at concentrations ranging from 0.5 - 10 µM and 0.1 - 5 µM respectively. Inhibitors were applied in the rinse buffer under arterial shear (1000 s^−1^). The formation, and subsequent treatment, of platelet aggregates was monitored on a brightfield microscope during each experiment (Supplementary Note 1). After completion of flow experiments, aggregates were fixed and stained for CD41 and F-actin and ExM-prepared to qualitatively determine effects of actin inhibitors on platelet morphology, platelet size and actin organisation (Supplementary Figure 3).

Under control conditions (Supplementary Figure 3A), platelets at the base of the aggregate had filopodia and lamellipodia-like protrusions and an interconnected F-actin network, whist the platelets were more rounded with cortical actin at the top of the aggregate, as shown previously in Fig 1. Aggregates treated with 0.5 µM CytD (Supplementary Figure 3B) showed a reduction in the presence of actin-based protrusions. At all higher concentrations (Supplementary Figure 3C-E) the actin-based protrusions were absent, with platelets adopting a homogeneously round morphology throughout the aggregates. At 5 and 10 µM, we observed a reduction in platelet size and CD41 distribution was altered, becoming more punctate and discontinuous than in controls. Aggregates treated with 0.1 µM LatA (Supplementary Fig 3F) exhibited a disruption to actin protrusion at the adhesion surface similarly to aggregates treated with CytD. At 1 µM LatA (Supplementary Figure 3G) we observed a dramatic disruption to actin organisation at the surface, with actin present as punctate accumulations as opposed to the meshwork observed in controls. At 3 and 5 µM this disruption in actin organisation was accompanied by a change of platelet shape and production of vesicle-like accumulations in the direction of flow (Supplementary Figure 3I-J, yellow arrows).

### 4.5 Quantification of structural disruptions induced by actin inhibitors on pre-formed platelet aggregates

We next performed a detailed quantitative characterisation of the effects of actin inhibitors across multiple healthy donors. We used 1 µM LatA and 2 µM CytD as these concentrations resulted in a significant observable disruption to the cytoskeleton and platelet morphology without causing complete breakdown of the aggregates. Expanded aggregates, treated with the inhibitors and labelled for CD41 and actin were imaged by epifluorescence and SPIM imaging (Supplementary Figure 4, Figure 5). Supplementary Videos 5-7 are supplied to highlight observed structural alterations under control and actin inhibited conditions.

**Figure 5.**
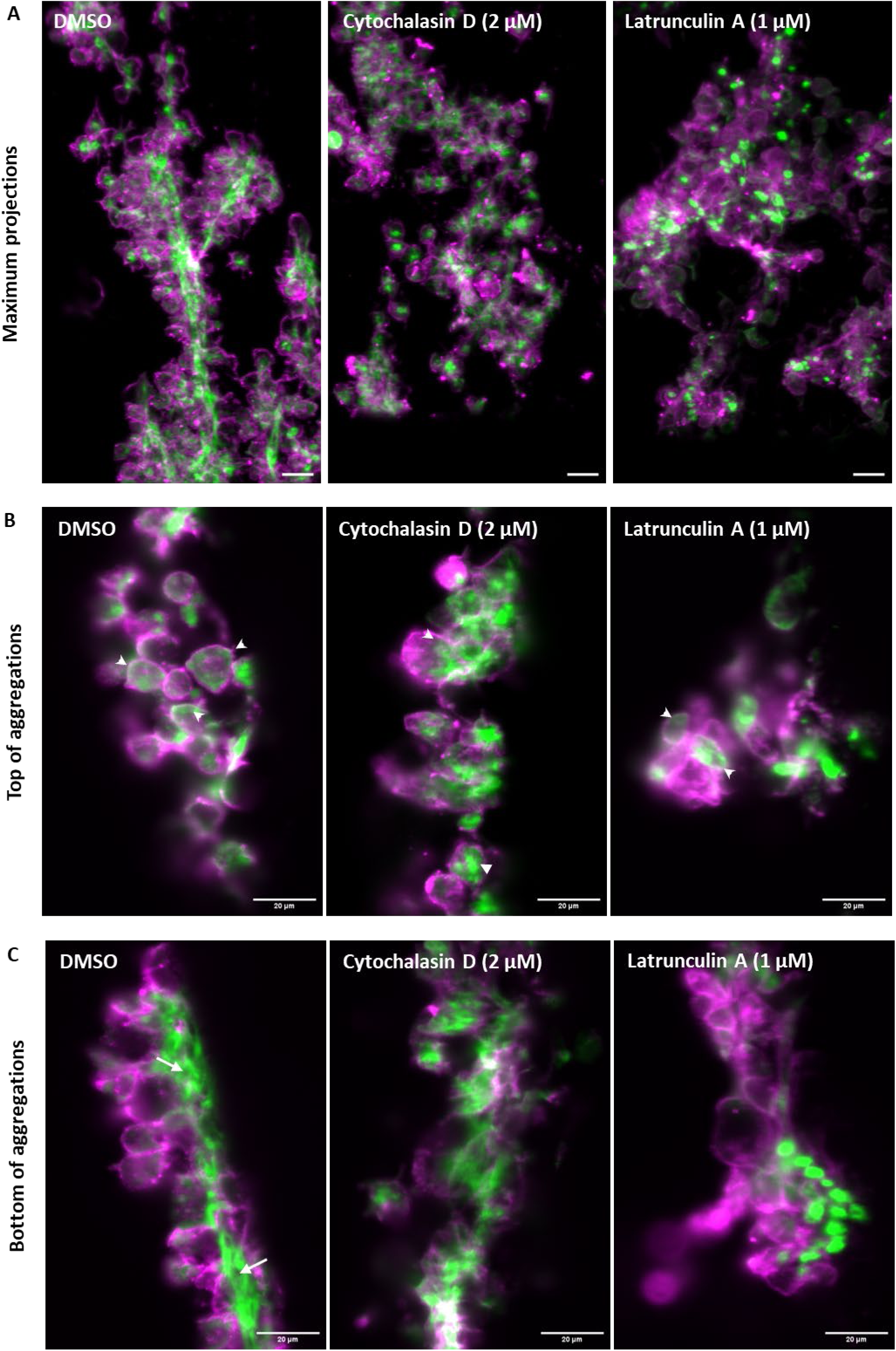
Characterising the structural impact of actin inhibitors on pre-formed platelet aggregates under arterial shear. Platelet aggregates were formed under arterial shear in a parallel plate flow chamber. CytD (2 µM) or LatA (1 µM) was flowed over aggregates for 10 minutes at arterial shear. Aggregates were fixed, immunostained (CD41 – magenta, F-actin – green) and prepared via ExM. Images were acquired on a SPIM. A) maximum projections. B) Example focal planes taken at the top of the aggregate or C) near the coverslip for each condition. Scale bars - 20 µm. White arrows – actin networks, white arrowheads – cortical actin. N=3.

Under control conditions, we observed platelets with lamellipodia- and filopodia-like actin protrusions spread across the adhesion surface, and a highly organised actin meshwork at the adhesion surface (Figure 5C, white arrows). By contrast, we observed rounded platelets with cortically organised actin at the top of the control aggregates (Figure 5B, white arrowheads). Intriguingly, this cortical organisation of actin close to the platelet membrane was retained in aggregates treated with both actin inhibitors (Figure 5B, white arrowheads). However, at the adhesion surface, the actin meshwork was completely absent following treatment with both actin inhibitors. Platelets appeared to cover reduced surface area and have changed in shape. Following treatment with LatA, we observed a total loss of the actin network, replaced by dense, punctate actin accumulations. These observations suggest actin may be more dynamic at the adhesion surface than at the top of the aggregates.

To quantitatively describe the structural alterations induced in platelet aggregates by actin inhibition, we applied the image analysis workflow as described previously. This revealed that the median platelet volume (Figure 6A(i)), boundary voxels (Figure 6B(i)), boundary : volume ratio (Figure 6C(i)) and sphericity (Figure 6D(i)) were all affected by F-actin disruption. Although significantly different, many of the changes are subtle and so plotting frequency distributions (Figure 6A(ii) – 6D(ii)) helps to demonstrate the changes. For example, the frequency distribution of the boundary : volume ratio of platelets treated with both CytD and LatA is left-shifted compared to controls (Figure 6C(ii)) indicating a smaller surface area to volume ratio. Whereas in Figure 6D, the LatA-treated platelets are right shifted, indicating a more spherical shape, demonstrating the role actin dynamics plays in shape maintenance.

**Figure 6.**
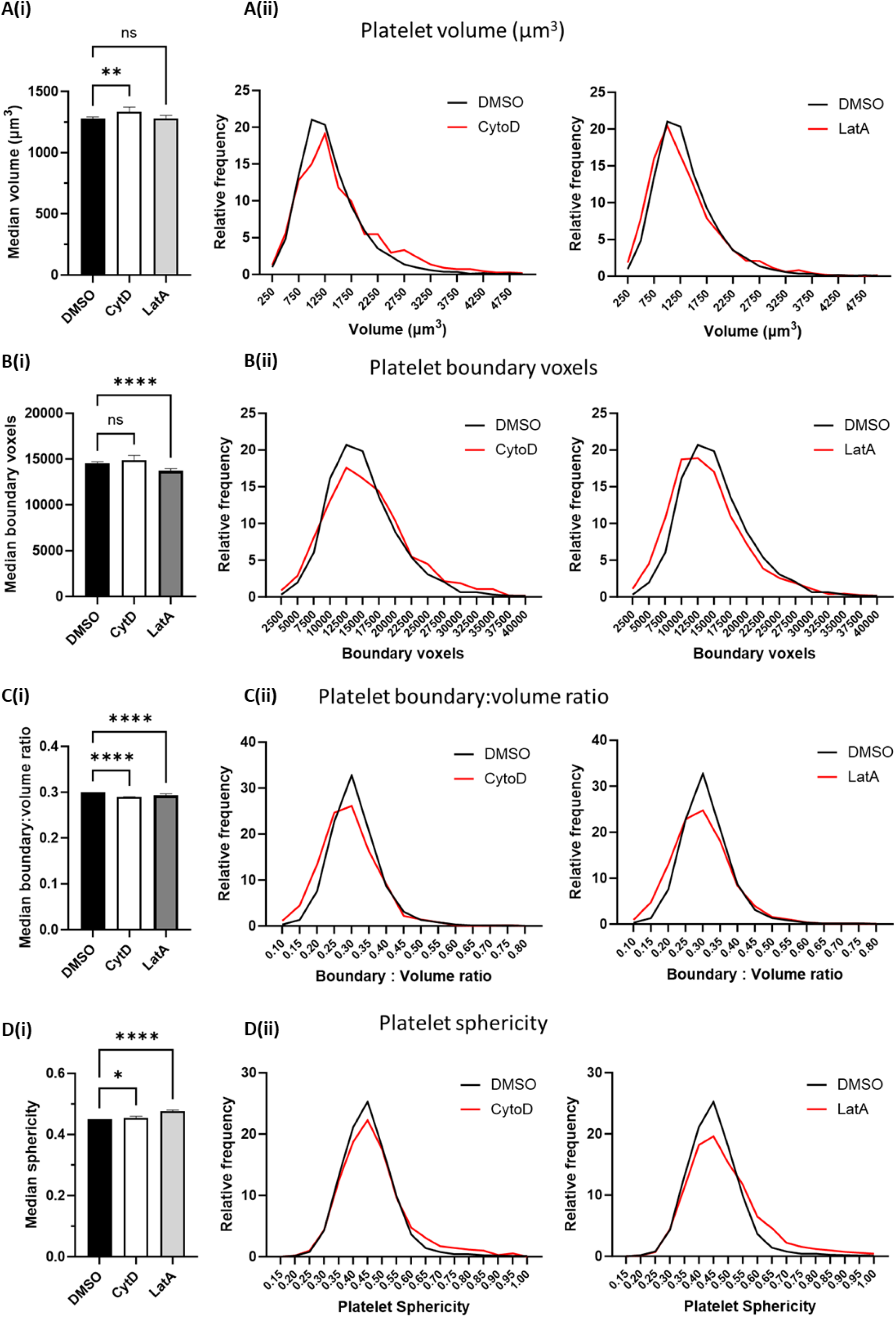
Quantification of the effect of actin disruption on morphology of pre-formed platelet aggregates. Platelet aggregates treated with either CytD (2 µM) or LatA (1 µM) were expanded and imaged via the workflow described in figure 3. Plots demonstrating the effect on A) platelet volume, B) Platelet boundary voxels, C) volume : boundary ratio and D) sphericity demonstrate the ability of the technique to identify subtle changes in platelet morphology. Bars in A(i), B(i), C(i) & D(i) represent median ± 95 % confidence interval. Number of platelets analysed; control, n = 4967; CytD, n = 1123; LatA, n = 4509. Asterix represent significance of treated samples compared to control (DMSO) established by Kruskal-Wallis and Dunns multiple comparison test. ns = not significant, * = P<0.05, ** = P<0.01, **** = P<0.0001. n = 3 replicates. A(ii), B(ii), C(ii) & D(ii) show the frequency distribution for each parameter in response to either CytD (left) or LatA (right).

Finally, the distribution of F-actin between membrane and cytoplasmic compartments was assessed after treatment with actin polymerisation inhibitors. Treatment with both inhibitors caused a significant reduction in the intensity of staining on both compartments (Figure 7A & B), as would be expected due to actin depolymerisation, as phalloidin only binds to F-actin. However, the data also confirmed the observation that there was a shift in the distribution of actin away from the membrane and towards the central regions, as indicated by a significant decrease in the membrane : cytoplasmic ratio (Figure 7C).

**Figure 7.**
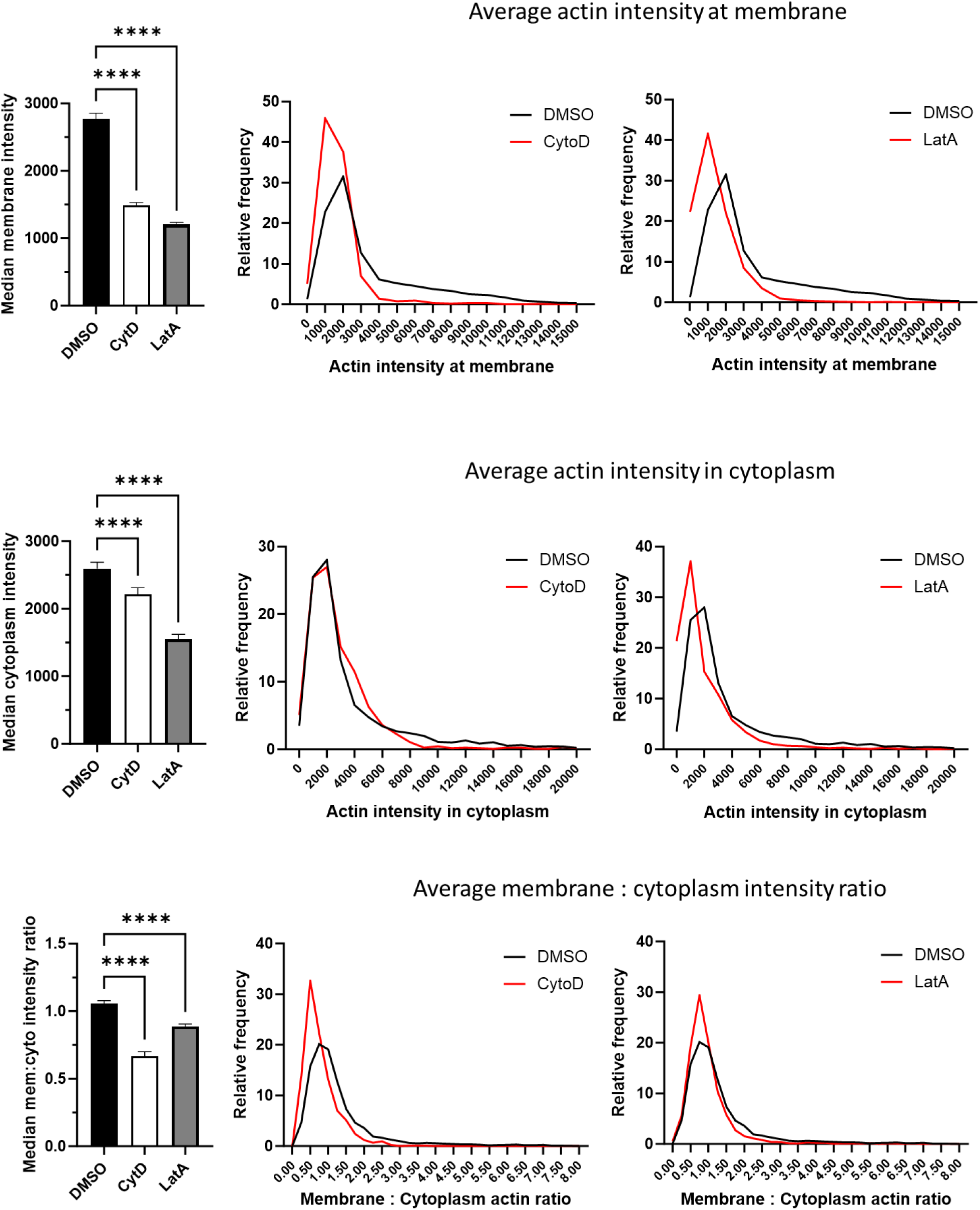
Quantification of the effect of actin disruption on F-actin distribution in pre-formed platelet aggregates. A) average intensity of the actin signal in membrane proximal voxels of segmented platelets. B) average intensity of the actin signal in cytoplasmic voxels of segmented platelets. C) average membrane : cytoplasmic ratio of F-actin signal. Bars in A(i), B(i) & C(i) represent median ± 95 % confidence interval. Number of platelets analysed; control, n = 4967; CytD, n = 1123; LatA, n = 4509. Asterix represent significance of treated samples compared to control (DMSO) established by Kruskal-Wallis and Dunns multiple comparison test. **** = P<0.0001. n = 3 replicates. A(ii), B(ii), C(ii) & D(ii) show the frequency distribution for each parameter in response to either CytD (left) or LatA (right).

## 5 Discussion

Thrombus formation is a complex process, resulting in heterogeneous organisation of components (e.g. platelets, neutrophils, fibrin) dependent on several parameters including type of vessel perturbation, shear rates and vascular site [13–15]. Developing a 3D model of platelet aggregation with nanoscale resolution will provide critical insight into their composition, mechanism of formation and factors contributing to thrombus resolution. Microscopy is an indispensable tool in characterising the architecture of haemostatic and thrombotic platelet aggregates [34, 35], allowing for high temporal (e.g. intravital microscopy) and spatial (e.g. electron microscopy) resolution imaging [16, 36]. Yet these approaches are not amenable to 3D volumetric imaging of whole platelet aggregates with nanoscale resolution. To overcome these limitations, we apply ExM to platelet-rich aggregates formed on collagen at arterial shear rate to qualitatively and quantitatively describe the heterogeneous structural organisation of membrane and cytoskeletal components within thrombi.

We applied epifluorescence and selective plane illumination microscopy (SPIM) on expanded platelet aggregates to produce a comprehensive dataset comprised of hundreds of platelet aggregates. The former approach is widely accessible and allows description of gross architecture of platelet aggregates. The latter approach offers (i) improved optical sectioning capabilities; (ii) high signal-to-noise ratio in each image slice; (iii) minimisation of optical aberrations due to the water immersion refractively matching the expanded specimens; (iv) rapid image acquisition [24, 37].

Initial inspection of 3D images of platelet aggregates labelled for platelet integrin, CD41, tubulin and F-actin revealed a heterogeneous distribution of platelets as discerned using CD41 as a membrane marker, and heterogeneous organisation of subcellular cytoskeletal components. At the basal surface, platelets possessed lamellipodia and filopodia-like actin protrusions with networks of F-actin bundles running through multiple platelets. By contrast, platelets closer to the top of aggregates adopted a round morphology with actin present cortically beneath the membrane. Our previous work identified the presence of actin nodules at the basal surface of platelets during early adhesion and spreading [3] and that platelets lacking actin nodule formation had reduced thrombus stability [4]. This led us to hypothesise that platelet actin nodules might form between platelets in thrombi to increase stability. However, in the current study, we have not observed formation of nodules within aggregates but have demonstrated the presence of extensive actin bundles throughout the thrombi. This is in line with observations by Schurr et. al. [38] and leads us to believe that these bundles are a major factor in thrombus stability.

Under perturbed conditions, where preformed platelet aggregates were treated with actin inhibitors CytD and LatA, we observed the networks of F-actin bundles at the basal surface were disrupted whilst round platelets with cortical actin at the apical surface were retained. The observed stress fibre-like F-actin bundles may be a result of platelets at the adhesion surface interacting with collagen and von Willebrand factor (vWF), a critical step in platelet adhesion. These actin-myosin cytoskeletal structure would impart mechanical strength and stability under arterial shear [39], as indicated by a loss of stability and structural organisation observed when actin inhibitors are applied. Interestingly, under control conditions these bundles appear to coincide with the collagen filament alignment on the coverslip, consistent with observations that collagen receptor, GPVI, aligns on collagen fibres in both spread and aggregated platelets [25, 40]. This actin filament alignment is absent following treatment with inhibitors, suggesting continuous actin remodelling at the adhesion surface. It is interesting to consider whether GPVI interactions with collagen, and actin filament bundle formation/organisation could be linked and contribute to thrombus stability. The reduced impact of actin inhibition in rounder, apical platelets where actin is cortically organised, may suggest the actin in these platelets is less dynamic.

To quantitatively describe structural alterations in platelet aggregates, achieved here by applying actin inhibitors, we developed a custom analysis and visualisation workflow to allow extraction of several aggregate features (e.g. platelet volume, sphericity). We also demonstrate a capability to describe the subcellular organisation of F-actin across populations of aggregates. Following actin inhibition, an overall reduction of F-actin in both the membrane and cytosolic regions was detected; this was more pronounced in LatA-treated aggregates. The membrane: cytosolic ratio of actin was also reduced; more so in the CytD-treated aggregates. These differences may be derived from the different mechanisms by which these compounds disrupt F-actin. CytD binds to F-actin filaments and prevents addition of actin monomers, initially stabilising filaments, but ultimately disrupting the equilibrium causing filament disassembly [41]. Conversely, LatA binds and sequesters actin monomers, preventing addition to the filament and causing rapid filament depolymerisation [42].

This study demonstrates ExM as a powerful technique, capturing 3D structural organisation of thousands of individual platelets within hundreds of platelet aggregates with relative ease and speed when compared with other super-resolution techniques (e.g. structured illumination microscopy (SIM) [43, 44]. ExM relies on expanding fixed specimens in an isotropic manner within a polyelectrolyte gel, resulting in a spreading out of features and improved resolution. An inherent property of this preparation is optical clearing, overcoming the optical opacity of thrombi which continues to hamper comprehensive visualisation and description of their architecture. Our combined custom analysis workflow is complementary, allowing the user to assess several parameters which can be interrogated and improve modelling of thrombus architecture. For example, given our observations of heterogeneity in platelet organisation throughout aggregates, we may analyse different pools of platelets. Platelet measurements could be separated based on sphericity, to account for flatter platelets at the basal surface versus rounder platelets at the apical side, thus more comprehensive analysis can be incorporated into our provided workflow.

Further we are not limited to platelets, by defining parameters for segmentation clearly, this approach would allow for investigation of multiple components (e.g. red blood cells (RBCs), fibrin). The ability to describe subcellular features may be extended to other features within platelets (e.g. α-granules). Correlating subcellular organisation of features with overall cellular or morphological features will powerfully allow the user a holistic overview of thrombus structure. We anticipate the workflow we have developed can be applied to patient-derived thrombi which has important implications in understanding thrombus formation, resolution and clinical intervention [45].

## Acknowledgements

The authors acknowledge the help of Dee Kavanagh and Joao Correia (COMPARE) in image acquisition and processing, Natalie Jooss (University of Birmingham) for training in parallel plate flow chamber experiments, Dirk-Peter Herten (University of Birmingham) for support during final data acquisition and paper submission. The authors also acknowledge the support of the Platelet Group at the University of Birmingham. This research was supported by the infrastructure provided by Advanced Research Computing at the University of Birmingham including data storage on the Research Data Store and computations on the BlueBEAR HPC service. This research has been funded by The British Heart Foundation (NH/18/3/33913, Accelerator AA/18/2/34218) and The Centre for Membrane Proteins and Receptors (COMPARE). The National Institute of Health and Care Research (NIHR) Birmingham Biomedical Research Centre (NIHR203326) has supported the University of Birmingham Institute of Cardiovascular Sciences where this research is based. The opinions expressed in this paper are those of the authors and do not represent any of the listed organizations.

## Authorship contributions

Conceptualization: E.L.F., S.G.T., N.S.P.; Methodology: E.L.F., J.A.P., E.G., S.G.T., N.S.P.; Software: J.A.P.; Validation: E.L.F.; Formal analysis: E.L.F., J.A.P., S.G.T.; Investigation: E.L.F., E.G., S.G.T., N.S.P.; Resources: N.S.P., S.G.T.; Data curation: E.L.F.; Writing - original draft: E.L.F., N.S.P., S.G.T.; Writing - review & editing: E.L.F., J.A.P., E.G., R.K.N., I.B.S., S.P.W., N.S.P., S.G.T.; Visualization: E.L.F., E.G.; Supervision: N.S.P., S.G.T.; Project administration: N.S.P., S.G.T.; Funding acquisition: J.A.P., R.K.N., I.B.S., S.P.W. N.S.P., S.G.T.

## References

1 Thomas SG. 3 - The Structure of Resting and Activated Platelets. In: Michelson AD, ed. Platelets (Fourth Edition): Academic Press, 2019, 47–77.

2 Bearer EL, Prakash JM, Li Z. Actin dynamics in platelets. Int Rev Cytol. 2002; 217: 137–82. 10.1016/s0074-7696(02)17014-8.

3 Calaminus SD, Thomas S, McCarty OJ, Machesky LM, Watson SP. Identification of a novel, actin-rich structure, the actin nodule, in the early stages of platelet spreading. J Thromb Haemost. 2008; 6: 1944–52. 10.1111/j.1538-7836.2008.03141.x.

4 Poulter NS, Pollitt AY, Davies A, Malinova D, Nash GB, Hannon MJ, Pikramenou Z, Rappoport JZ, Hartwig JH, Owen DM, Thrasher AJ, Watson SP, Thomas SG. Platelet actin nodules are podosome-like structures dependent on Wiskott-Aldrich syndrome protein and ARP2/3 complex. Nat Commun. 2015; 6: 7254. 10.1038/ncomms8254.

5 Janus-Bell E, Mangin PH. The relative importance of platelet integrins in hemostasis, thrombosis and beyond. Haematologica. 2023; 108: 1734–47. 10.3324/haematol.2022.282136.

6 Pollitt AY, Grygielska B, Leblond B, Désiré L, Eble JA, Watson SP. Phosphorylation of CLEC-2 is dependent on lipid rafts, actin polymerization, secondary mediators, and Rac. Blood. 2010; 115: 2938–46. 10.1182/blood-2009-12-257212.

7 Poulter NS, Pollitt AY, Owen DM, Gardiner EE, Andrews RK, Shimizu H, Ishikawa D, Bihan D, Farndale RW, Moroi M, Watson SP, Jung SM. Clustering of glycoprotein VI (GPVI) dimers upon adhesion to collagen as a mechanism to regulate GPVI signaling in platelets. J Thromb Haemost. 2017; 15: 549–64. 10.1111/jth.13613.

8 Atkinson L, Yusuf MZ, Aburima A, Ahmed Y, Thomas SG, Naseem KM, Calaminus SDJ. Reversal of stress fibre formation by Nitric Oxide mediated RhoA inhibition leads to reduction in the height of preformed thrombi. Sci Rep. 2018; 8: 3032. 10.1038/s41598-018-21167-6.

9 Calaminus SD, Auger JM, McCarty OJ, Wakelam MJ, Machesky LM, Watson SP. MyosinIIa contractility is required for maintenance of platelet structure during spreading on collagen and contributes to thrombus stability. J Thromb Haemost. 2007; 5: 2136–45. 10.1111/j.1538-7836.2007.02696.x.

10 Yusuf MZ, Raslan Z, Atkinson L, Aburima A, Thomas SG, Naseem KM, Calaminus SDJ. Prostacyclin reverses platelet stress fibre formation causing platelet aggregate instability. Sci Rep. 2017; 7: 5582. 10.1038/s41598-017-05817-9.

11 Bender M, Palankar R. Platelet Shape Changes during Thrombus Formation: Role of Actin-Based Protrusions. Hamostaseologie. 2021; 41: 14–21. 10.1055/a-1325-0993.

12 Huang J, Li X, Shi X, Zhu M, Wang J, Huang S, Huang X, Wang H, Li L, Deng H, Zhou Y, Mao J, Long Z, Ma Z, Ye W, Pan J, Xi X, Jin J. Platelet integrin αIIbβ3: signal transduction, regulation, and its therapeutic targeting. J Hematol Oncol. 2019; 12: 26. 10.1186/s13045-019-0709-6.

13 Alkarithi G, Duval C, Shi Y, Macrae FL, Ariens RAS. Thrombus Structural Composition in Cardiovascular Disease. Arterioscler Thromb Vasc Biol. 2021; 41: 2370–83. 10.1161/ATVBAHA.120.315754.

14 Hashimoto T, Kunieda T, Honda T, Scalzo F, Sharma LK, Hinman JD, Rao NM, Nour M, Bahr-Hosseini M, Saver JL, Raychev R, Liebeskind DS. Heterogeneity between proximal and distal aspects of occlusive thrombi on pretreatment imaging in acute ischemic stroke. Neuroradiol J. 2022; 35: 378–87. 10.1177/19714009211049713.

15 Staessens S, De Meyer SF. Thrombus heterogeneity in ischemic stroke. Platelets. 2021; 32: 331–9. 10.1080/09537104.2020.1748586.

16 Stalker TJ, Traxler EA, Wu J, Wannemacher KM, Cermignano SL, Voronov R, Diamond SL, Brass LF. Hierarchical organization in the hemostatic response and its relationship to the platelet-signaling network. Blood. 2013; 121: 1875–85. 10.1182/blood-2012-09-457739.

17 Tomaiuolo M, Matzko CN, Poventud-Fuentes I, Weisel JW, Brass LF, Stalker TJ. Interrelationships between structure and function during the hemostatic response to injury. Proc Natl Acad Sci U S A. 2019; 116: 2243–52. 10.1073/pnas.1813642116.

18 Schrottmaier WC, Assinger A. The Concept of Thromboinflammation. Hamostaseologie. 2024; 44: 21–30. 10.1055/a-2178-6491.

19 Weisel JW, Litvinov RI. Visualizing thrombosis to improve thrombus resolution. Res Pract Thromb Haemost. 2021; 5: 38–50. 10.1002/rth2.12469.

20 Chernysh IN, Nagaswami C, Kosolapova S, Peshkova AD, Cuker A, Cines DB, Cambor CL, Litvinov RI, Weisel JW. The distinctive structure and composition of arterial and venous thrombi and pulmonary emboli. Sci Rep. 2020; 10: 5112. 10.1038/s41598-020-59526-x.

21 Jolugbo P, Ariens RAS. Thrombus Composition and Efficacy of Thrombolysis and Thrombectomy in Acute Ischemic Stroke. Stroke. 2021; 52: 1131–42. 10.1161/STROKEAHA.120.032810.

22 Chozinski TJ, Halpern AR, Okawa H, Kim HJ, Tremel GJ, Wong RO, Vaughan JC. Expansion microscopy with conventional antibodies and fluorescent proteins. Nat Methods. 2016; 13: 485–8. 10.1038/nmeth.3833.

23 Faulkner EL, Thomas SG, Neely RK. An introduction to the methodology of expansion microscopy. The International Journal of Biochemistry & Cell Biology. 2020; 124: 105764. 10.1016/j.biocel.2020.105764.

24 Gao R, Asano SM, Boyden ES. Q&A: Expansion microscopy. BMC Biology. 2017; 15: 50. 10.1186/s12915-017-0393-3.

25 Jooss NJ, Smith CW, Slater A, Montague SJ, Di Y, O’Shea C, Thomas MR, Henskens YMC, Heemskerk JWM, Watson SP, Poulter NS. Anti-GPVI nanobody blocks collagen- and atherosclerotic plaque-induced GPVI clustering, signaling, and thrombus formation. J Thromb Haemost. 2022; 20: 2617–31. 10.1111/jth.15836.

26 Pearce AC, Senis YA, Billadeau DD, Turner M, Watson SP, Vigorito E. Vav1 and vav3 have critical but redundant roles in mediating platelet activation by collagen. J Biol Chem. 2004; 279: 53955–62. 10.1074/jbc.M410355200.

27 Van Kruchten R, Cosemans JM, Heemskerk JW. Measurement of whole blood thrombus formation using parallel-plate flow chambers - a practical guide. Platelets. 2012; 23: 229–42. 10.3109/09537104.2011.630848.

28 Schindelin J, Arganda-Carreras I, Frise E, Kaynig V, Longair M, Pietzsch T, Preibisch S, Rueden C, Saalfeld S, Schmid B, Tinevez J-Y, White DJ, Hartenstein V, Eliceiri K, Tomancak P, Cardona A. Fiji: an open-source platform for biological-image analysis. Nature Methods. 2012; 9: 676–82. 10.1038/nmeth.2019.

29 van der Walt S, Schönberger JL, Nunez-Iglesias J, Boulogne F, Warner JD, Yager N, Gouillart E, Yu T. scikit-image: image processing in Python. PeerJ. 2014; 2: e453. 10.7717/peerj.453.

30 Kluyver T, Ragan-Kelley B, Pérez F, Granger BE, Bussonnier M, Frederic J, Kelley K, Hamrick JB, Grout J, Corlay S, Ivanov P, Avila D, Abdalla S, Willing C, Team JD. Jupyter Notebooks - a publishing format for reproducible computational workflows. International Conference on Electronic Publishing, 2016.

31 Tillberg PW, Chen F, Piatkevich KD, Zhao Y, Yu CC, English BP, Gao L, Martorell A, Suk HJ, Yoshida F, DeGennaro EM, Roossien DH, Gong G, Seneviratne U, Tannenbaum SR, Desimone R, Cai D, Boyden ES. Protein-retention expansion microscopy of cells and tissues labeled using standard fluorescent proteins and antibodies. Nat Biotechnol. 2016; 34: 987–92. 10.1038/nbt.3625.

32 Stringer C, Wang T, Michaelos M, Pachitariu M. Cellpose: a generalist algorithm for cellular segmentation. Nature Methods. 2021; 18: 100–6. 10.1038/s41592-020-01018-x.

33 Auger JM, Watson SP. Dynamic tyrosine kinase-regulated signaling and actin polymerisation mediate aggregate stability under shear. Arterioscler Thromb Vasc Biol. 2008; 28: 1499–504. 10.1161/atvbaha.108.167296.

34 de Witt SM, Swieringa F, Cavill R, Lamers MM, van Kruchten R, Mastenbroek T, Baaten C, Coort S, Pugh N, Schulz A, Scharrer I, Jurk K, Zieger B, Clemetson KJ, Farndale RW, Heemskerk JW, Cosemans JM. Identification of platelet function defects by multi-parameter assessment of thrombus formation. Nat Commun. 2014; 5: 4257. 10.1038/ncomms5257.

35 Staessens S, François O, Desender L, Vanacker P, Dewaele T, Sciot R, Vanhoorelbeke K, Andersson T, De Meyer SF. Detailed histological analysis of a thrombectomy-resistant ischemic stroke thrombus: a case report. Thromb J. 2021; 19: 11. 10.1186/s12959-021-00262-1.

36 Brass LF, Tomaiuolo M, Welsh J, Poventud-Fuentes I, Zhu L, Diamond SL, Stalker TJ. 20 - Hemostatic Thrombus Formation in Flowing Blood. In: Michelson AD, ed. Platelets (Fourth Edition): Academic Press, 2019, 371–91.

37 Huisken J, Stainier DY. Selective plane illumination microscopy techniques in developmental biology. Development. 2009; 136: 1963–75. 10.1242/dev.022426.

38 Schurr Y, Sperr A, Volz J, Beck S, Reil L, Kusch C, Eiring P, Bryson S, Sauer M, Nieswandt B, Machesky L, Bender M. Platelet lamellipodium formation is not required for thrombus formation and stability. Blood. 2019; 134: 2318–29. 10.1182/blood.2019002105.

39 Atkinson L, Yusuf MZ, Aburima A, Ahmed Y, Thomas SG, Naseem KM, Calaminus SDJ. Reversal of stress fibre formation by Nitric Oxide mediated RhoA inhibition leads to reduction in the height of preformed thrombi. Scientific Reports. 2018; 8: 3032. 10.1038/s41598-018-21167-6.

40 Pallini C, Pike JA, O’Shea C, Andrews RK, Gardiner EE, Watson SP, Poulter NS. Immobilized collagen prevents shedding and induces sustained GPVI clustering and signaling in platelets. Platelets. 2021; 32: 59–73. 10.1080/09537104.2020.1849607.

41 Schliwa M. Action of cytochalasin D on cytoskeletal networks. J Cell Biol. 1982; 92: 79–91. 10.1083/jcb.92.1.79.

42 Spector I, Shochet NR, Blasberger D, Kashman Y. Latrunculins--novel marine macrolides that disrupt microfilament organization and affect cell growth: I. Comparison with cytochalasin D. Cell Motil Cytoskeleton. 1989; 13: 127–44. 10.1002/cm.970130302.

43 Knight AE, Gomez K, Cutler DF. Super-resolution microscopy in the diagnosis of platelet granule disorders. Expert Rev Hematol. 2017; 10: 375–81. 10.1080/17474086.2017.1315302.

44 Pluthero FG, Kahr WHA. Evaluation of human platelet granules by structured illumination laser fluorescence microscopy. Platelets. 2023; 34: 2157808. 10.1080/09537104.2022.2157808.

45 Zhao Y, Bucur O, Irshad H, Chen F, Weins A, Stancu AL, Oh EY, DiStasio M, Torous V, Glass B, Stillman IE, Schnitt SJ, Beck AH, Boyden ES. Nanoscale imaging of clinical specimens using pathology-optimized expansion microscopy. Nat Biotechnol. 2017; 35: 757–64. 10.1038/nbt.3892.

