## Supplemental information for "Expansion microscopy allows quantitative characterisation of structural organisation of platelet aggregates"

For

**List of Supplementary information included**

**1. Supplementary notes**

Supplementary note 1 – determining optimal concentration of latrunculin A and cytochalasin D to disrupt pre-formed platelet aggregates.

**2. Supplementary Figures**

Supplementary figure 1 - ExM reveals heterogeneous distribution of membrane and cytoskeletal components in platelet aggregates.

Supplementary figure 2 - Donor variation in platelet aggregates.

Supplementary figure 3 - Treatment of pre-formed platelet aggregates with Cytochalasin D and Latrunculin A under flow results in structural changes.

Supplementary figure 4 - Treatment of pre-formed platelet aggregates with Cytochalasin D and Latrunculin A under flow results in structural changes.

**3. Supplementary Tables**

Supplementary Table 1 – Summary of identified platelets and platelet aggregates associated with Data Supplement 1 and Data Supplement 2.

**4. Supplementary Files**

Data Supplement 1 – Data Explorer – Donor experiments

Data Supplement 2 – Data Explorer – Actin inhibitor experiments

**5. Supplementary Videos**

Supplementary Videos 1 - 7

**Supplementary Note 1**

Optimal doses of latrunculin A and cytochalasin D were determined by applying each drug at a range of concentrations (0.1 - 5 μM and 0.5 - 10 μM respectively). Under control conditions, we observed low numbers of single platelets embolising in the direction of flow during treatment.

In the aggregates treated with latrunculin A, groups of platelets embolised in the direction of flow within 3 minutes during the rinse at 1, 2, 3 and 5 μM. Embolisation of platelets was markedly less at 0.1 μM latrunculin A. At 3 and 5 μM concentrations, we further noted platelets losing their round morphology and spreading in the direction of flow.

In the aggregates treated with cytochalasin D, we observed embolization of single platelets in the direction of flow during the rinse at 0.5, 2, 5 and 10 μM. The embolization of platelets was much lower in the presence of cytochalasin D than latrunculin A.

### Supplementary Figures

#### Supplementary Figure 1

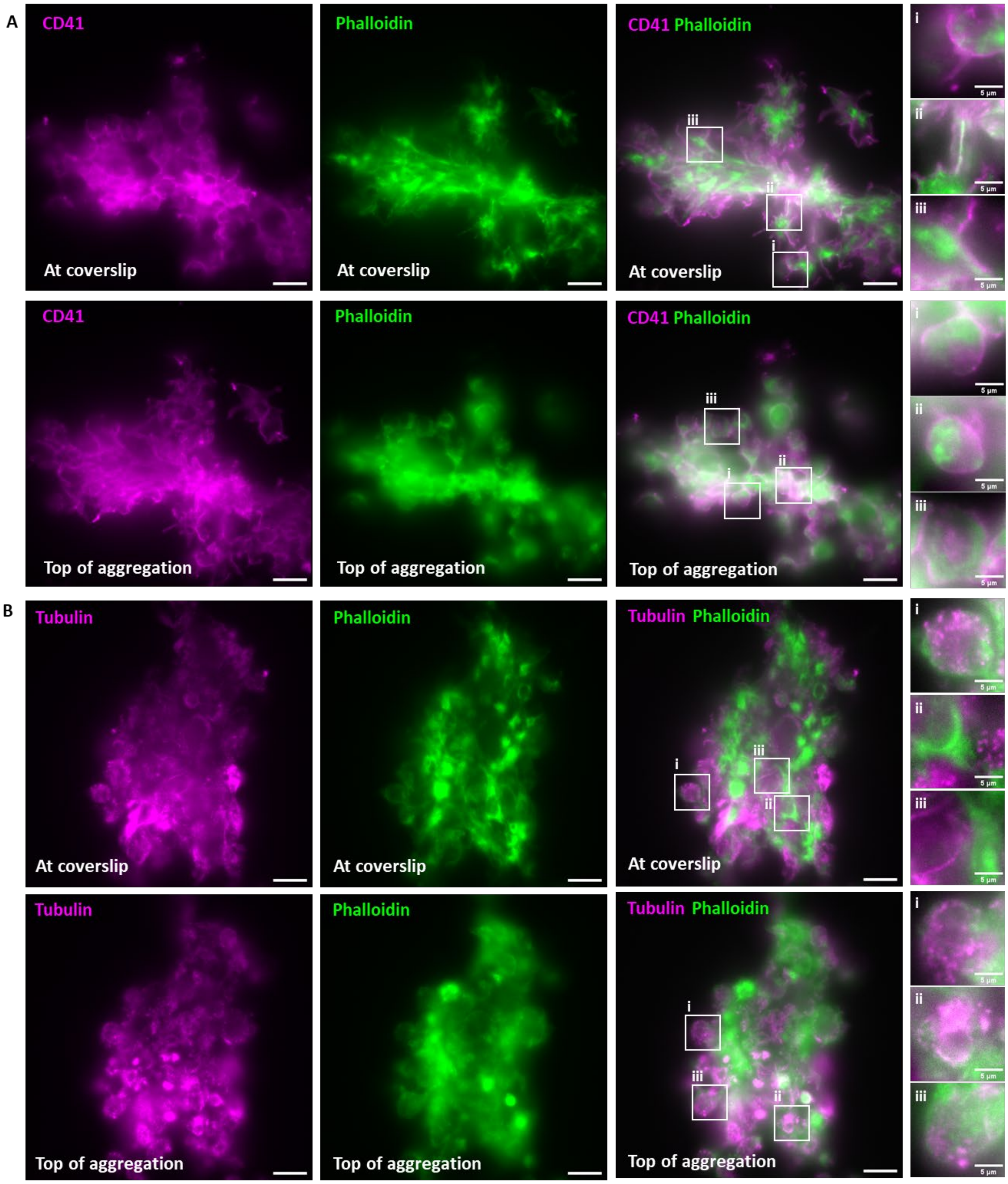

**Supplementary Figure 1. Expansion microscopy (ExM) reveals heterogeneous distribution of membrane and cytoskeletal components in platelet aggregates.** Platelet aggregates were prepared in parallel plate flow chambers under arterial shear (1000 $s^{-1}$) rate on collagen-coated coverslips. Aggregates were fixed, immunostained and expanded before imaging by widefield epifluorescence microscopy. A) Example images of aggregates stained for CD41 (magenta) and F-actin (green) are shown. Slices at the coverslip (upper panel) and at the top of the aggregate (lower panel) are provided. Scale bars 20 μm (inset image scale bars 5 μm). B) Images of aggregates stained for tubulin (magenta) and F-actin (green) are shown. Slices at the coverslip (upper panel) and at the top of the aggregate (lower panel) are provided. Inset images show features of interest. Scale bars 20 μm (inset image scale bars 5 μm). n=5.

### Supplementary Figure 2

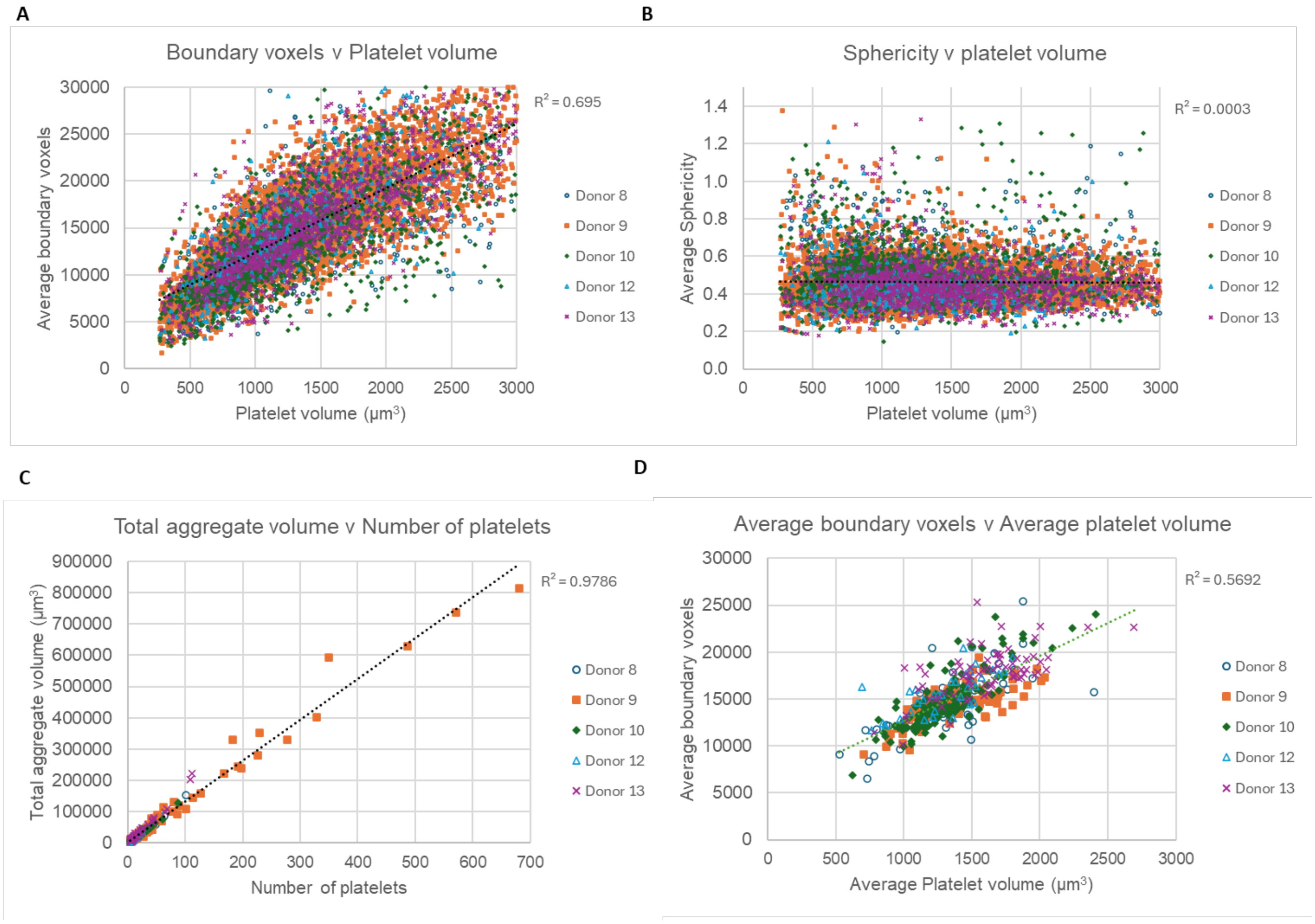

**Supplementary Figure 2. Example data plots from image analysis workflows.** Scatter plots of A) platelet boundary voxels versus platelet volume, and B) platelet sphericity versus platelet volume on all individually segmented platelets from 5 different donors. Scatter plots of C) total aggregate volume versus number of platelets per aggregate and H) average boundary voxels versus average platelet volume from aggregates identified from 5 different donors by Otsu thresholding. Full data sets with the ability to filter plots by donor or measured parameters are provided in Data Supplement 1.

#### Supplementary Figure 3

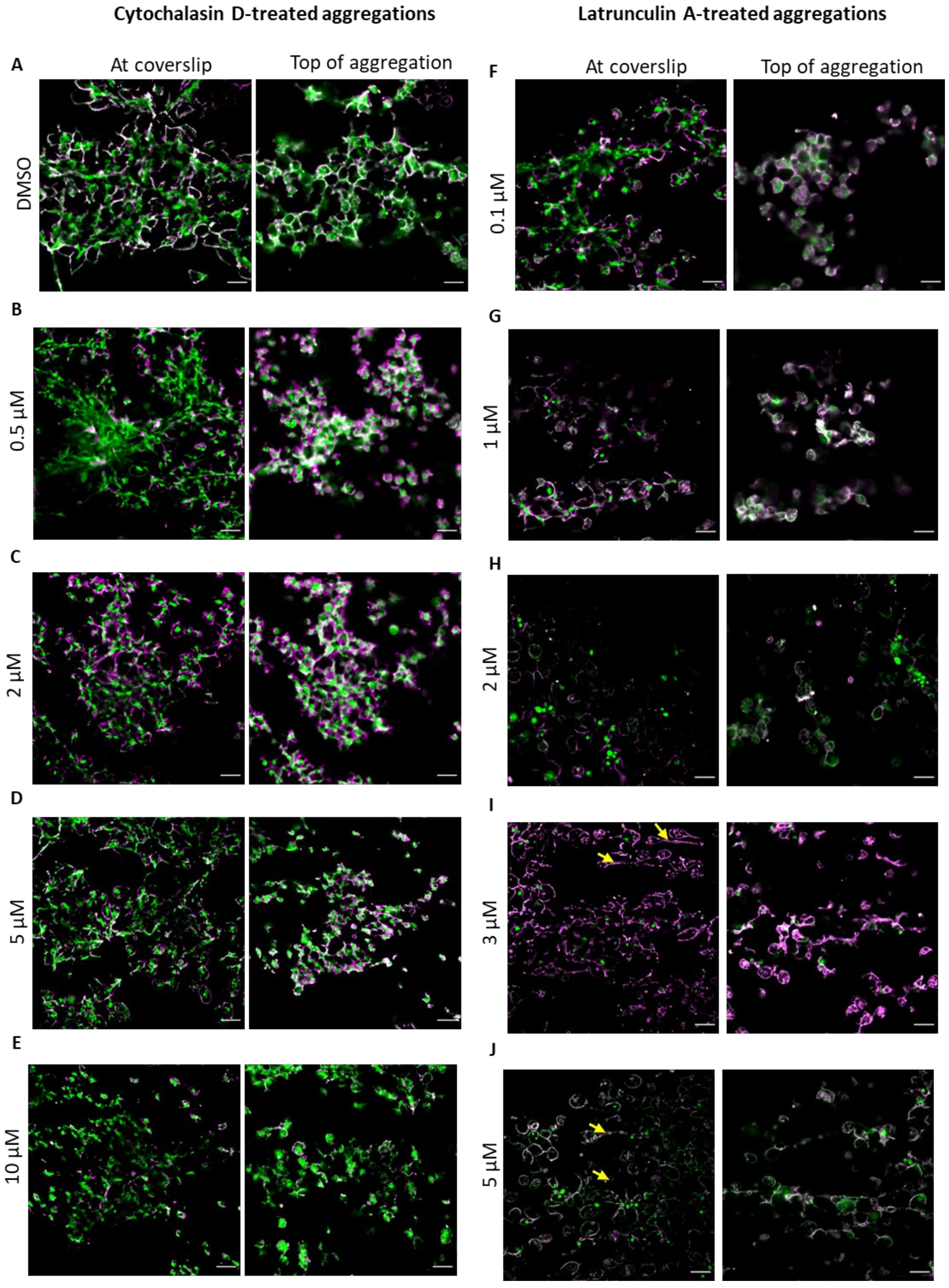

**Supplementary Figure 3. Treatment of pre-formed platelet aggregates with Cytochalasin D and Latrunculin A under flow results in structural changes.** Platelet aggregates were formed in parallel plate flow chambers on coverslips coated with collagen. Pre-formed aggregates were treated at the stated concentrations of actin inhibitor for 10 minutes under flow. Fixed aggregates were stained for CD41 (magenta) and actin (green). ExM was completed and then post-expansion images acquired on an epifluorescence microscope and deconvolved. Scale bars 20 μm. n=3.

**Supplementary Figure 4**

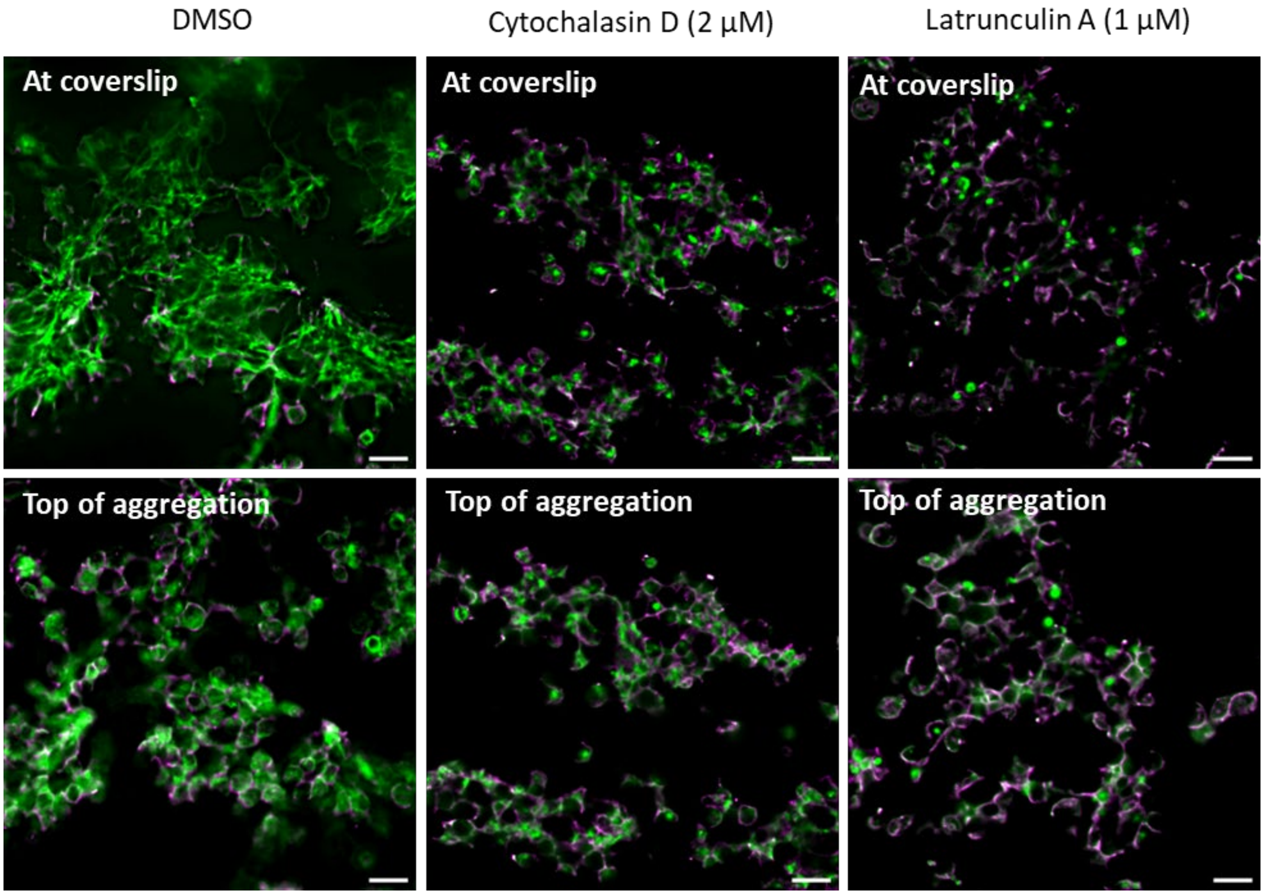

**Supplementary Figure 4. Treatment of pre-formed platelet aggregates with Cytochalasin D and Latrunculin A under flow results in structural changes.** Platelet aggregates were formed in parallel plate flow chambers on coverslips coated with collagen. Pre-formed aggregates were treated at the stated concentrations by diluting treatment in rinse buffer and incubating for 10 minutes under flow. Fixed aggregates were stained for CD41 (magenta) and actin (green). ExM was completed and then post-expansion images acquired on an epifluorescence microscope and deconvolved. Scale bars 20 μm. Representative images from n=7.

#### Supplementary Tables

##### Supplementary Table 1

| Experiment design | Number of identified platelet aggregates | Number of individual platelets |
|---|---|---|
| Platelet aggregates formed from five healthy donors as shown in Figure 2A-B and 3. | **421** | **11,471** |
| Platelet aggregates formed from healthy donors and then treated with either: DMSO, Cytochalasin D or Latrunculin A as shown in Figure 5-7. | **329** | **10,879** |

#### Data Supplements

2 Microsoft Excel workbooks are provided as data supplements. These contain all the raw data generated from the python scripts for the donor (Supplement 1) and actin inhibitor (Supplement 2) experiments. (Note – the raw data tabs are hidden for simplicity).

In each workbook are 3 tabs

*Individual Platelet Data tab* – this contains the measured values of volume (voxels, and $\mu m^3$), boundary voxels, sphericity and boundary : volume ratio for each identified platelet. The data slicer buttons can be used to filter this data by donor, image series and identified aggregate. Note the data is pre-filtered to a minimum object volume of 268 $\mu m^3$ – this is to filter out any very small objects (i.e. less than the volume of a resting platelet) from the analysis.

*Aggregate Summary Data tab* – this contains the summary statistics of total aggregate volume ($\mu m^3$), number of platelet per aggregate, average platelet volume ($\mu m^3$), average boundary voxels, average sphericity and average boundary : volume ratio for each identified aggregate. The data slicer buttons can be used to filter this data by donor. Note the data is pre-filtered to a minimum object volume of 268 $\mu m^3$ – this is to filter out any very small objects (i.e. less than the volume of a resting platelet) from the analysis.

*Data Explorer tab* – this contains summary tables and example plots for a range of outputs. The left-hand side is drawn from the *Individual Platelet Data* tab and the right-hand side the *Aggregate Summary Data* tab. The plotted data will change depending on the data slicer filter applied in the other tabs, or it can be filtered directly using the data slicer buttons in this tab.

**Supplementary Videos**

Supplementary Video 1 – Platelet aggregates formed under arterial shear stained for CD41 (magenta) and F-actin (green) and prepared via fourfold expansion microscopy protocol. Video of slice-by-slice acquisition on selective plane illumination microscope in stage scanning mode.

Supplementary Video 2 – Platelet aggregates prepared as in Supplementary Video 1. Acquired images on selective plane illumination microscope were deskewed and presented as 3D projections rotating along the y-axis.

Supplementary Video 3 - Platelet aggregates formed under arterial shear stained for tubulin (magenta) and F-actin (green) and prepared via fourfold expansion microscopy protocol. Video of slice-by-slice acquisition on selective plane illumination microscope in stage scanning mode.

Supplementary Video 4 - Platelet aggregates prepared as in Supplementary Video 2. Acquired images on selective plane illumination microscope were deskewed and presented as 3D projections rotating along the y-axis.

Supplementary Video 5 – Platelet aggregates formed under arterial shear were treated for 10 minutes with dimethylsulfoxide (0.001% (v/v) in rinse buffer under shear. Following 2 minutes of stasis, platelet aggregates were rinsed for 2 minutes and then fixed. Aggregates were then stained for CD41 (magenta) and F-actin (green) and prepared via fourfold expansion microscopy protocol. Video of slice-by-slice acquisition on selective plane illumination microscope in stage scanning mode.

Supplementary Video 6 - Platelet aggregates formed under arterial shear were treated for 10 minutes with 2 µM cytochalasin D in rinse buffer under shear. Following 2 minutes of stasis, platelet aggregates were rinsed for 2 minutes and then fixed. Aggregates were then stained for CD41 (magenta) and F-actin (green) and prepared via fourfold expansion microscopy protocol. Video of slice-by-slice acquisition on selective plane illumination microscope in stage scanning mode.

Supplementary Video 7 - Platelet aggregates formed under arterial shear were treated for 10 minutes with 1 µM latrunculin A in rinse buffer under shear. Following 2 minutes of stasis, platelet aggregates were rinsed for 2 minutes and then fixed. Aggregates were then stained for CD41 (magenta) and F-actin (green) and prepared via fourfold expansion microscopy protocol. Video of slice-by-slice acquisition on selective plane illumination microscope in stage scanning mode.
